# Towards a multi-metric assessment of river restoration ecological outcomes across projects and contexts

**DOI:** 10.64898/2025.12.10.693400

**Authors:** Blandine Charrat, Mathieu Floury, Martin Palt, Alienor Jeliazkov, Stefan Stoll, Evelyne Tales, Ralf C.M. Verdonschot, Christine Weber, Armin Lorenz, Céline Le Pichon, Jérémy Piffady

## Abstract

The assessment of the ecological outcomes of river restoration projects remains a complex undertaking due to the high variability of responses observed. This variability is dependent on a number of factors, including the location of the site under study, the indicators and taxa utilised in the analysis. In this study, a multi-metric and multi-contextual approach was applied to analyse 62 river restoration projects located in France, Germany and the Netherlands. This methodological approach facilitated the characterisation of the responses exhibited by macroinvertebrate communities in response to restoration initiatives, and the subsequent identification of the environmental contexts and the categories of restoration measures that exerted influence on these responses. Utilising log-response ratios based on both taxonomic and functional ecological metrics, a k-means clustering analysis was performed. This analysis yielded two distinct groups of sites: the first exhibiting predominantly positive responses to restoration, and the second demonstrating neutral or marginally negative responses. Positive responses have been shown to be associated with ambitious restoration projects, which are characterised by measures targeting regional-scale processes. These measures include the implementation of projects designed to improve river longitudinal continuity in degraded catchment areas, where the density of obstacles and the intensity of agricultural use are particularly high. Conversely, restoration initiatives targeting local-scale issues within more natural contexts or at high altitude exhibited minimal to no ecological response. This study emphasises the significance of an integrated approach that incorporates various facets (taxonomic and functional) of biodiversity, along with a meticulous examination of the environmental context and project construction, to enhance the evaluation of the ecological benefits of river restoration. The findings of this study may also be applicable in the context of fieldwork, providing practitioners with a framework to inform the design of restoration projects.

## Introduction

River hydromorphological restoration projects are being implemented across the globe with the primary objective of enhancing the ecological status of watercourses that have been degraded due to various pressures, particularly those of anthropogenic origin (Reid et al. 2019; Dudgeon et al. 2006). As ecosystem functioning is directly linked to this ecological status, and as the costs of those measures are often very high, it is therefore essential to understand how the systems react to those restoration measures and whether the restoration projects benefit the site’s ecological status. This is of particular importance in the context of biodiversity, as it provides a framework for guiding water management decisions. In recent decades, there has been a concerted effort to study the species richness and diversity patterns exhibited by rivers that have undergone restoration measures (Sinclair et al. 2023). The restoration effects have been shown to vary significantly between taxa in different studies (Palmer et al. 2010; Schmutz et al. 2014; Pilotto et al. 2019). Many case studies have documented a wide range of outcomes, including positive, neutral, and negative results.

The varying outcomes observed in ecological restoration initiatives can be primarily attributed to the metrics employed for the assessment of biodiversity, which exhibit significant variability across studies. It is evident that studies are distinguished by disparate objectives, resulting in the utilisation of varied metrics. Consequently, a comparison of their respective results is rendered complex. For instance, functional metrics (based on species traits) have been shown to exhibit stronger effects of restoration than taxonomic metrics (based on species identities) (Al-Zankana et al. 2020; Kail et al. 2015), although the inverse has also been observed (England and Wilkes 2018; Seidel et al. 2021).). However, it should be noted that the effects may vary even among functional metrics. This phenomenon can be illustrated by examining the current velocity preferences exhibited by fish species. Specifically, rheophilic species appear to derive greater benefit from restoration initiatives compared to limnophilic species (Mueller et al. 2014; Stoll et al. 2014).

Furthermore, numerous factors have been demonstrated to exert influence on restoration outcomes. These factors can be linked to two main areas. Firstly, the environment, including both natural and anthropogenic conditions of the pre-restored site. Secondly, the implementation of the restoration project itself, including project design and implemented measures. The variability in terms of natural environmental conditions and characteristics of anthropogenic pressures between restored sites may also influence the ecological effects of restoration projects. For instance, it appears crucial to consider the natural gradients. Indeed, as Manfrin et al. (Manfrin et al. 2019) demonstrate, sites at higher altitude tend to have a longer response time to restoration than sites at lower altitude. Furthermore, the pre-restoration condition, in terms of land use types in the catchment in interaction with its geo-climatic conditions, can greatly influence restoration outcomes (Feld et al. 2016). The construction of the restoration project is another source of variability in restoration outcomes. For instance, the length of the restored section has been demonstrated to influence the efficacy of the restoration project for fish in a positive manner (Schmutz et al. 2014). Despite the fact that Hering et al. (2015) did not identify this effect in their study on multiple groups of aquatic and floodplain-inhabiting organisms, they concluded that sections shorter than 2 km are inefficient for biodiversity recovery, a conclusion that is in line with a meta-analysis carried out by (Kail et al. 2015) for aquatic organisms. Furthermore, the outcomes of restoration endeavours may be subject to variation in accordance with the nature of the restoration measures employed on the degraded site. Instream measures have been demonstrated in several studies to yield more favourable outcomes in comparison to other types of measures, including riparian or channel planform measures (Al-Zankana et al. 2020; Kail et al. 2015). These outcomes have been observed in terms of taxonomic, functional abundance, richness, and diversity. Furthermore, measures that enhance connectivity are expected to promote species diversity by facilitating the movement of individuals from source populations to restored sections (England et al. 2021). A plethora of studies have investigated one or several of these effects, either at the level of individual restoration projects, or with comparative analyses based on monitoring data from multiple restoration projects, or with meta-analyses on published studies. Nevertheless, to the best of our knowledge, no integrative study combining all of these metrics and factors together has been conducted to evaluate their combined influence on restoration outcomes. Such a study would appear to be a significant improvement, as it would provide a comprehensive picture.

The initial phase in the conceptualisation of a methodology for the evaluation of outcomes in river restoration initiatives might entail the implementation of a multi-metric approach. This would facilitate the identification of outcomes patterns, with consideration given to the various biodiversity expressions articulated through the diverse ecological metrics. The subsequent stage of the research will entail the identification of the impact of the environmental context and the restoration project itself on the various outcome patterns that emerged in the initial phase of the study.

The objective of this study is to ascertain the principal factors that influence river biodiversity responses to hydromorphological river restoration. In order to achieve this objective, it is proposed that a cross-project, multi-metric and multi-contextual approach be adopted for the study of the ecological response of macroinvertebrate communities to river restoration. Macroinvertebrates are considered to be pivotal taxa in the context of restoration, as they are frequently employed as bioindicators of freshwater ecosystem health (Bonada et al. 2006; Mondy et al. 2012; Prat et al. 2009) and are presumed to exhibit rapid responses to restoration measures (Paillex et al. 2009; Simons et al. 2001).

The objective of the present study is to utilise a clustering approach for the purpose of grouping restored sites based on the mean ecological responses of their macroinvertebrate communities. In a subsequent step, the environmental context or restoration measure characteristics that may drive the observed partition among sites is determined. To the best of our knowledge, this approach has never been used in the context of ecological responses to river restoration. Examples can be found in studies of ecosystem services (Raudsepp-Hearne et al. 2010; Renard et al. 2015).

In light of the variability observed in the extant literature and the multiplicity of response metrics under study, it is predicted that different response patterns will emerge. Therefore, the dataset will be divided into multiple clusters, with each cluster consisting of sites that demonstrate analogous restoration outcomes. Secondly, it is predicted that utilising this multi-metric and cross-project approach will facilitate the identification of favourable contexts for river restoration. Consequently, this will enable the drawing of conclusions to guide practitioners in selecting the most suitable combination of sites and restoration measures.

## Material and Methods

Figure 1 provides a synopsis of the statistical methodologies employed in the present study. All analyses were conducted utilising R version 4.3.3 (R Core Team 2024).

**Figure 1:**
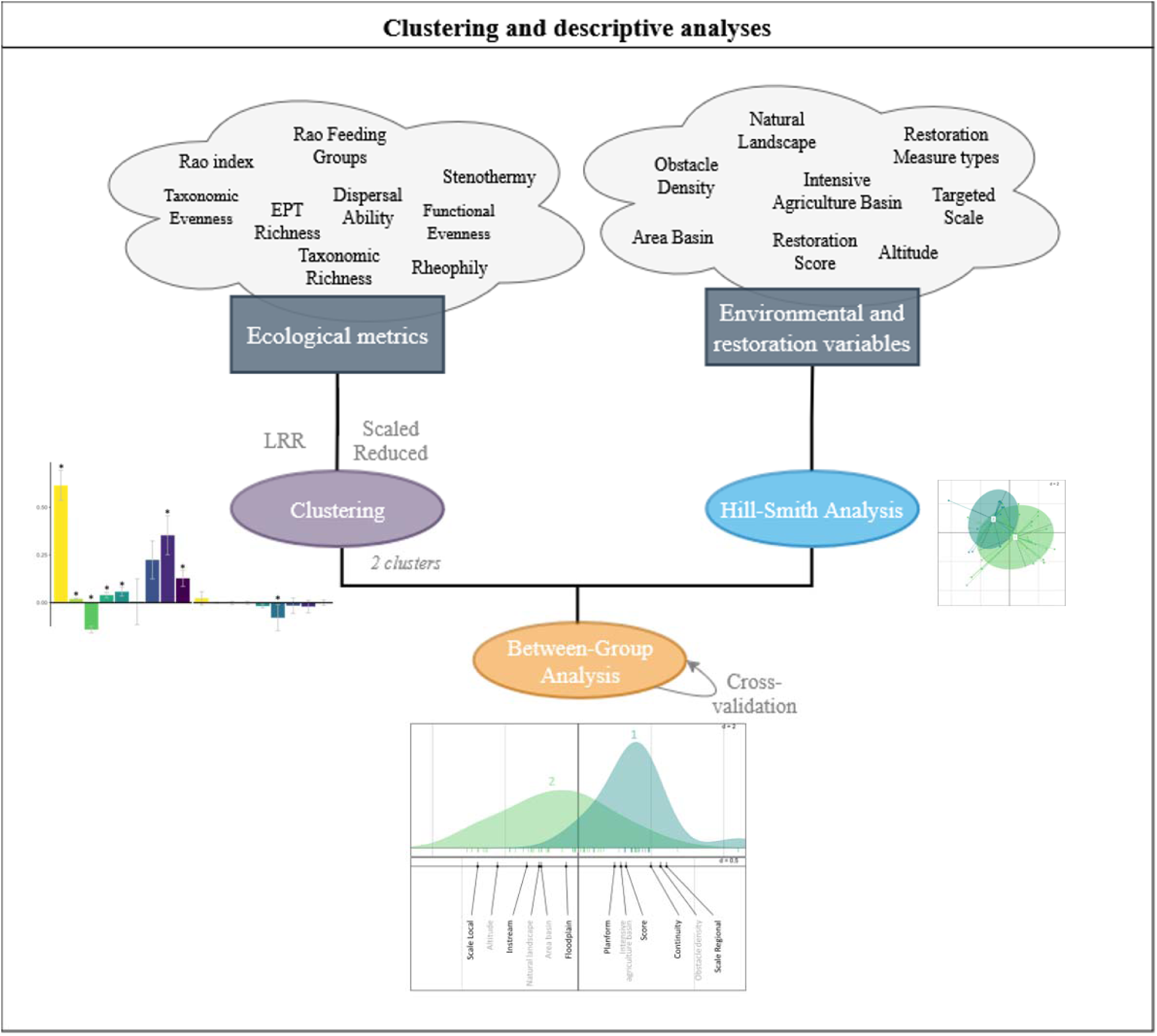
Summary of the analyse processes for clustering and descriptive analyses led in the study. LRR = log-response ratios.

### 1. Data preparation

The present study employs a total of 9 ecological metrics, adopting a multi-metric approach with a particular emphasis on sensitive taxa and dispersal abilities. Furthermore, given that the dataset under consideration comprises 62 sites which vary with respect to environmental factors, geographical location and the implementation of restoration measures, the approach adopted is multi-contextual.

The primary objective of this study is to calculate the response metrics to restoration.

#### 1.a. Calculation of response metrics to restoration

A total of 62 sites were surveyed, distributed across France (n = 26), Germany (n = 9), and the Netherlands (n = 27), where reach-scale restoration measures had been implemented between 1988 and 2018 (see Figure 2 for details). The measures implemented were intended to enhance the following aspects of fluvial morphology: instream, planform, floodplain, and river continuity. The effects of the fish were monitored using benthic macroinvertebrates, which were sampled in accordance with national standardised protocols. Furthermore, studies have shown that sites which have undergone restoration are frequently subject to rapid ecological succession in the initial years following the implementation of such measures. This has the effect of rendering it difficult to distinguish the proper effect of the restoration (Lepori et al. 2005). Consequently, the present study exclusively focused on the post-restoration sampling dates, which ranged from three to six years following the implementation of the measures. This timeframe was deemed sufficient for communities to stabilise following the potential ‘disturbances’ caused by restoration, particularly in regard to the short life cycle of the majority of macroinvertebrates. Taxonomic resolution exhibited variation primarily from species to family, yet remained constant over the monitoring period at a given site, culminating in a collection of 1000 taxa. The standardisation of collected taxa was conducted as densities per square metre.

**Figure 2:**
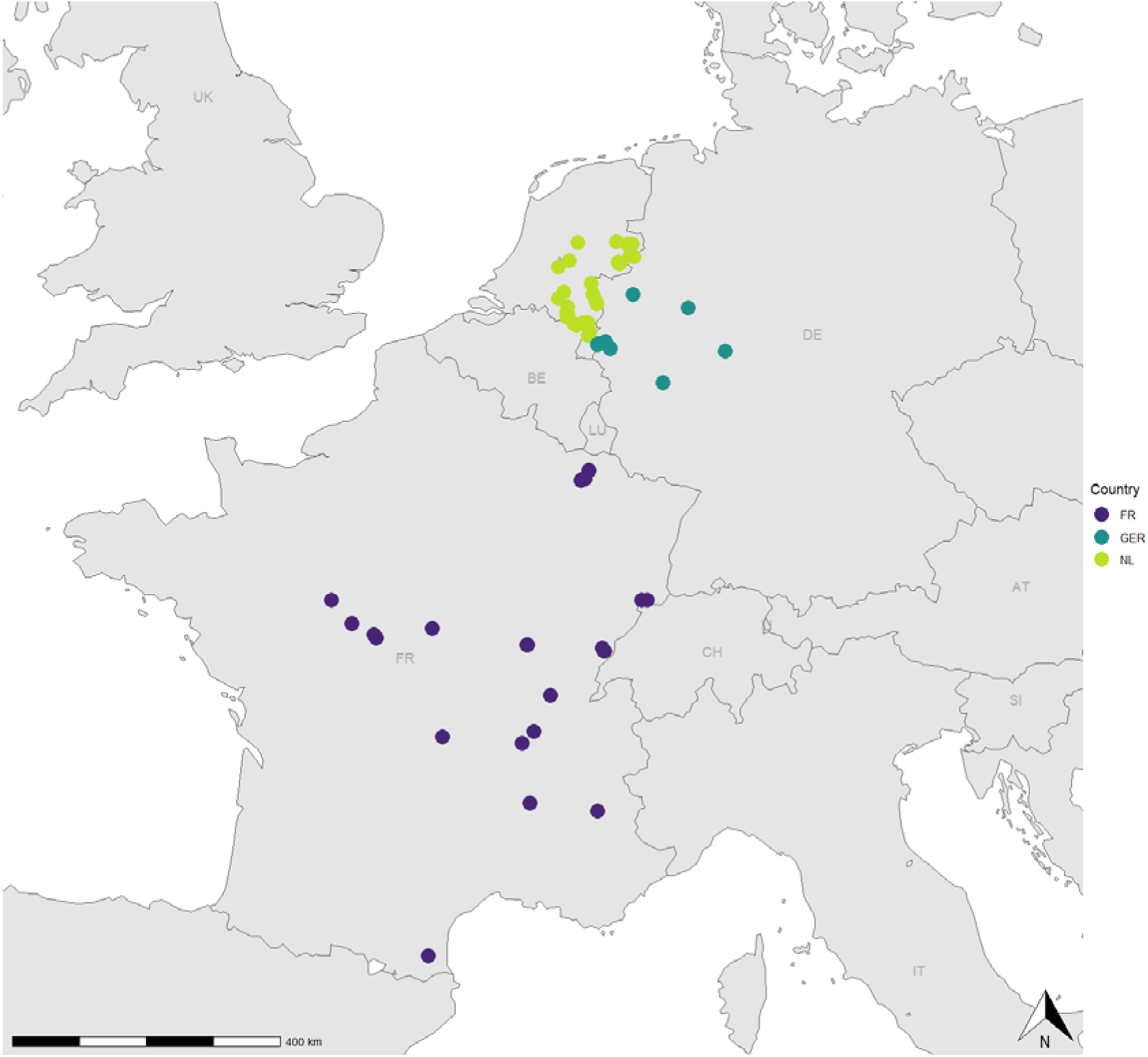
Location of the 62 restored sites studied across Europe.

We subsequently proceeded to define a series of taxonomy- and trait-based metrics with the objective of evaluating alterations in the richness and diversity of macroinvertebrate communities. Initially, the taxon richness (number of taxa) and evenness (taxon distribution) were calculated for each sample. The relative richness of the Ephemeroptera, Plecoptera and Trichoptera taxa (the so-called ‘EPT’ taxa, considered as sensitive taxa) in the communities was also calculated. Secondly, a functional space was defined on the basis of the following traits: (i) habitat preferences, (ii) dispersal and mobility, (iii) organism size and form, (iv) life history and reproduction, and (v) feeding groups (see Appendix 1: Table S1). The space was then used to predict which traits would respond following restoration (Pilotto et al. 2019).

The distribution of taxa in this multidimensional trait space was weighted according to their relative ln[x+1]-transformed density, and the Rao’s quadratic entropy (functional diversity) and the functional evenness were then computed using the *dbFD*() function from the FD package (Laliberté, Legendre, Shipley 2009). In a similar manner, the feeding group diversity (Rao’s entropy) was calculated on the basis of the functional subtypes defined using solely the ‘feeding habit’ trait.

Finally, an evaluation was conducted to ascertain the extent to which rheophily and cold-water stenothermy, in addition to the dispersal ability of the communities, might potentially serve as indicators of response to restoration. In order to accomplish this objective, individual scores of rheophily and stenothermy were defined for each taxon based on their fuzzy-coded affinities (i.e. scores between 0 and 5) for the trait modalities ‘fast’ (trait ‘current velocity’) and ‘psychrophylic’ (trait ‘temperature’), respectively. In accordance with the methodology established by Bonada et al. (Bonada et al. 2012), the individual scores of dispersal ability for each taxon were defined as the sum of their fuzzy-coded affinities for the modalities ‘aquatic passive’, ‘aquatic active’, ‘aerial passive’ and ‘aerial active’ (trait ‘dispersal mode’), respectively, weighted by 1, 5, 10 and 20. The standardisation of the three indices was achieved through the implementation of the maximum-minimum rescaling approach, thereby ensuring that the resulting individual scores ranged from 0 to 1. Subsequently, the community-weighted means (CWM) of rheophily, stenothermy and dispersal ability were determined by multiplying the individual scores of all taxa present in the community by their respective relative densities (e.g. Lavorel et al. 2008).

In order to assess changes in the different metrics in a standardised and comparable way, the effect sizes of the restoration were calculated using the Log-Response Ratio (LRR, Osenberg, Sarnelle, et Cooper 1997). A LRR was computed for each metric at each restored site and post-restoration sampling date, according to the monitoring design (‘Before-After’ [BA, 36% of the sites], ‘Control-Impact’ [CI, 30% of the sites] or full ‘Before-After-Control-Impact’ [BACI, 33% of the sites]):

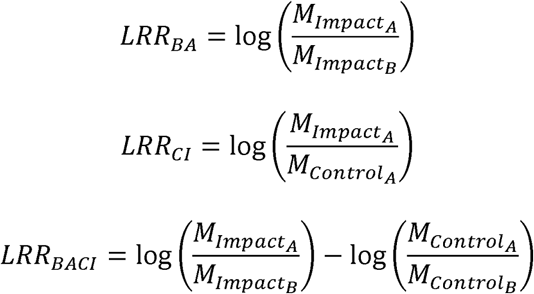

where M is a given metric, Control and Impact a pair of, respectively, unrestored and restored sites, and B and A, respectively, pre- and post-restoration years.

The LRR, when considered dimensionless, facilitates the comparison of responses based on disparate metrics (potentially in different units) and divergent monitoring designs, thereby ensuring an unbiased comparison. The presence of positive and negative values, respectively, signifies the occurrence of increases and decreases of the metric post-restoration. The LRRs were then averaged for each site over the different post-restoration dates within the 3-6-year period.

#### 1.b. Environmental context and restoration measure characteristics

In the context of the environmental study, the altitude (m) of the median points of the restored reaches was extracted from the MERIT digital elevation model (Yamazaki et al. 2017). The surface area of the drainage basins (km²) upstream of these median points was also calculated based on the WISE WFD database (European Environment Agency 2023). The additional anthropogenic pressures were described at different scales. On the one hand, the proportion of intensive agriculture across the upstream drainage basin, as well as the proportion of (near-)natural areas across the landscape corridor (i.e. 3-km long from 0.5 km downstream to 2.5 km upstream of the restored site, 100-m wide on both banks) were calculated using Corine Land Cover (European Environment Agency 2019). Conversely, the fragmentation of rivers upstream of the restoration sites was gauged by the density of obstacles, defined as the number of barriers per kilometre of river. In accordance with the AMBER Barrier Atlas (Belletti et al. 2020), all barriers were taken into consideration over the entire river network branching within a 10-kilometre "instream" isodistance upstream of the median points of the restored reaches. The processing of all spatial-context variables was conducted utilising the QGIS desktop version 3.22.14. The mean and standard error values are presented in Table 1.

**Table 1:** Environmental context and restoration measures characteristics of the studied sites. For restoration variables, “0” means that no measure aims at improving the concerned subject was led (e.g. continuity), “1” means that at least one measure aims at improving the concerned subject. The "Targeted Scale" variable gives a simplified view of the spatial scale of influence targeted by the restoration measure, with "local" (low impact) measures usually targeting small-scale processes and modifying up to 5km of river stretch (e.g. instream boulder additions) vs. “regional” (high impact) measures usually targeting larger-scale processes and modifying up to 20km of river stretch (e.g. river longitudinal and/or lateral connectivity restorations).

| *Environmental variable* | *Mean* | *Sd* |
| --- | --- | --- |
| Natural landscape (% of buffer occupancy) | 21 | 28 |
| Intensive agriculture basin (% of buffer occupancy) | 40 | 25 |
| Altitude (m) | 189 | 276 |
| Area basin (km<sup>2</sup>) | 209 | 312 |
| Obstacle density (number of structures on the river) | 0.67 | 0.8 |
| *Restoration variable* | *0* | *1* |
| Continuity | 39% | 61% |
| Instream | 50% | 50% |
| Floodplain | 73% | 27% |
| Planform | 21% | 79% |
|  | *Regional* | *Local* |
| Targeted scale restoration | 53% | 47% |

In terms of the restoration characteristics, the following factors were given full consideration: firstly, the targeted spatial scale of the restoration project (whether local or regional); secondly, the type(s) of measure(s) applied; and thirdly, a ‘score’ indicating the number of different measures applied within the same time frame during the restoration project. The ‘Targeted Scale’ variable is intended to provide a simplified representation of the spatial scale of influence targeted by the restoration measure. ‘Local’ measures, which typically stimulate small-scale processes (e.g. instream boulder additions), are characterised by a low spatial impact. In contrast, ‘regional’ measures, which generally aim to modify processes at larger scales (e.g. river longitudinal and/or lateral connectivity restorations), are characterised by a high spatial impact and are able to modify up to 20 km of river stretch. It is acknowledged that a single restoration initiative may encompass multiple measures, with each measure targeting distinct structures or processes (e.g. river continuity and/or morphology). In order to evaluate the correlation between the nature of restoration measures and the site clustering, a variable designated as "Restoration type" was determined based on an adapted Morandi’s classification (Morandi et al. 2017). The study encompassed four distinct methodologies, each designed to address a specific aspect of hydromorphological restoration, with the objective of enhancing the respective features of the target environment. The terms employed in this study are as follows: ‘continuity’ (river longitudinal continuity), ‘floodplain’ (floodplain morphology and lateral continuity), ‘instream’ (habitat heterogeneity) and ‘planform’ (form and geometry of the channel). As these methodologies are not mutually exclusive for a given restoration project, the factor was dummy-recorded by splitting it into the four methodologies and using ones (and zeroes) to indicate which restoration measures were applied (or not) at each site. Table 1 provides the proportion of sites for each type of measure and targeted scale of restoration. The variable "Score" was calculated, ranging from 1 when only one type of restoration measure was applied during the restoration work to 4 when the project incorporated the four measure types.

### Statistical analyses

#### 2.a. Identifying restoration response patterns: clustering

Given the sensitivity of clustering to high inter-metric variability, the average LRRs were scaled and centred to ensure the comparability of metric responses. The k-means clustering method, utilising the kmeans() function from the stats package, was then employed to ascertain the clusters according to these LRR values. Euclidian distance is the foundation of the k-means clustering algorithm. This algorithm aims to partition a dataset into k groups (= clusters) such that objects in one cluster are as similar as possible to each other and as dissimilar as possible to objects in other clusters. The fundamental principle underpinning this approach is to define clusters in a manner that ensures the total intra-cluster variation (Wtot) is minimised. The dataset under consideration meets the necessary conditions for the application of k-means clustering, as it is of sufficient size and the data has been standardised, thus eliminating any potential outliers. The optimal number of clusters was determined by employing the average silhouette method (Rousseeuw 1987) on the basis of the dataset (*fviz*_*nbclust*() function from factoextra package, Kassambara et Mundt 2020).

#### 2.b. Influence of environmental context and restoration measure characteristics: between-group analysis

Following the distribution of the sites into clusters, an investigation was conducted into the potential explanatory value of context characteristics with respect to both the environmental context and restoration characteristics of the sites on their association with each cluster. This investigation sought to elucidate the pattern with which the sites responded to restoration.

In order to appraise the influence of environmental context and restoration characteristics on the clustering, i.e. on restoration outcomes, a between-group analysis (BGA, Doledec & Chessel, 1987, 1989) was employed to obtain a descriptive analysis that maximises the variation of environmental context and restoration characteristics between clusters. In order to do so, firstly the Hill-Smith analysis (*dudi.hillsmith*() function of ade4 package, Dray et Dufour 2007), was run, on which the BGA was then applied with the *bca*() function of ade4 package. The BGA is responsible for determining the optimal axis that delineates the clusters within the space calculated by Hill-Smith analysis. The cluster identifier (1 or 2) of each site was employed as a group variable. BGA is an approach that permits the analysis of small sample sizes. This is due to the fact that it is based on the mean across sampling dates (between three and six years) of each variable describing the environmental context and restoration characteristics for each site (Doledec and Chessel 1987; 1989). This approach avoids potential problems linked to small samples. Furthermore, to ensure that our representation reflects reality, we implemented the cross-validation procedure proposed by the *randtest*() function, as outlined by Thioulouse et al. (2021). This procedure is contingent on the calculation of the Δ*O*□ij index (*loocv()* function, Thioulouse et al. 2021), which characterizes the percentage of difference between our representation of the BGA and the representations of cross-validated data. The lower this index is, the more the BGA representation is credible. The absolute contribution of explained variation for each variable in the BGA was calculated using the *inertia*.*dudi*() function (ade4 package, Dray et Dufour 2007).

## Results

### Identifying restoration response patterns

The application of the average silhouettes method to the clustering outputs indicated that the optimal number of clusters for the restored sites was two (see average silhouettes method representation in Appendix 2: Figure S1). The two clusters were distinguished by the average direction of the responses of the metrics. The initial cluster, comprising 16 sites, encompassed restoration initiatives that exhibited favourable outcomes across the majority of metrics, with the exception of functional evenness. The second cluster (comprising 46 sites) comprised restoration projects in which all metrics did not respond or responded negatively to restoration (see Figure 3).

**Figure 3:**
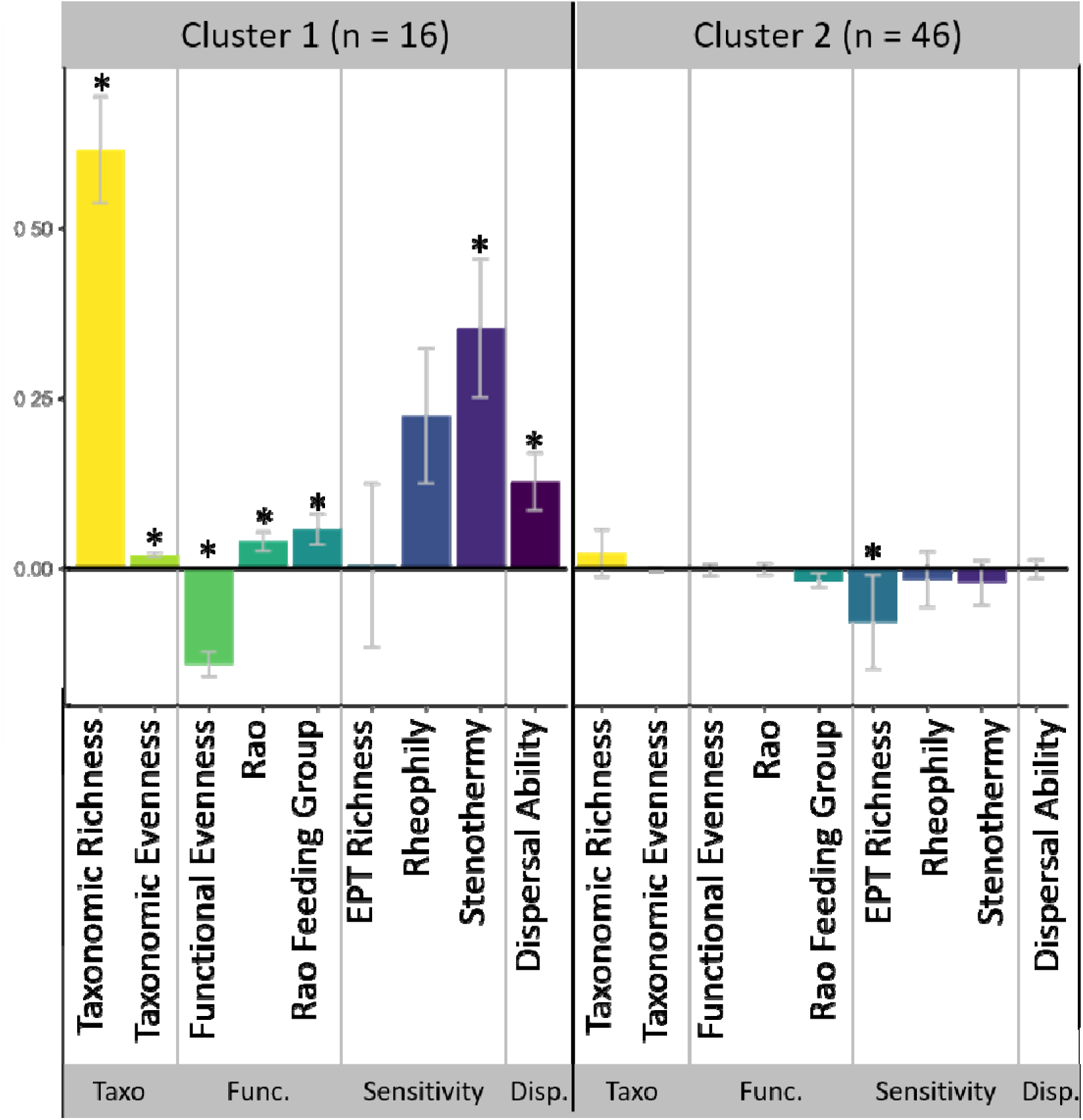
Average responses of ecological metrics (= average LRR) in the two clusters for macroinvertebrates. The height of the bar in the plot represents the mean value of the log response ratio, i.e. the response of each ecological metric. Here we present raw data and not scaled and centred data used for clusters determination, raw data allowing easier ecological interpretation. * indicates that the mean of the bar is different from 0. Taxo. = taxonomic metrics, Func. = functional metrics, Sensitivity = sensitivity metrics, Disp. = Dispersal ability metric. Error bars represent the standard error for each mean.

The strongest positive responses in cluster 1 were observed for taxonomic richness and for the community-weighted mean for rheophily and stenothermy. In the aforementioned sites, the dispersal ability metric has been observed to increase following restoration. Furthermore, functional diversity (Rao) and diversity in feeding groups (Rao Feeding Group) also increase. It is evident that communities occurring in sites of the cluster 1 are characterised by increased richness, sensitivity and functional diversity following restoration, thereby aligning with the established expectations. The majority of sites (cluster 2 = 46 out of 62) exhibited no response, or a negative response, to restoration measures when all metrics were considered collectively.

### Influence of environmental context and restoration measure characteristics

In the subsequent phase of the study, the investigation focused on the impact of environmental settings and characteristics of the restoration projects on the outcomes of the restoration.

The Between-Group Analysis (BGA) determined the axis that optimally differentiates the two groups in the Hill-Smith analysis (results presented in Appendix 3: Figure S1). The BGA has confirmed that the two clusters of sites, which are based on ecological responses, could be divided according to different environmental contexts and restoration characteristics. Indeed, the BGA representation (Figure 4) indicates that a division of the two clusters according to the environmental and restoration variables of the sites is possible. The representation of the BGA comprises a single axis, given that there are merely two clusters of restoration projects. The permutation test is found to be significant (p = 0.001), and the Δ*O*□ij index is determined to be null (0%), indicating that the BGA representation is credible. The two clusters accounted for 7.6% of the total variation. The restoration variables contributed to 60% of the explained variation, while the environmental variables contributed to 40% of the explained variation. The absolute contributions of each variable are presented in Figure 4.

**Figure 4:**
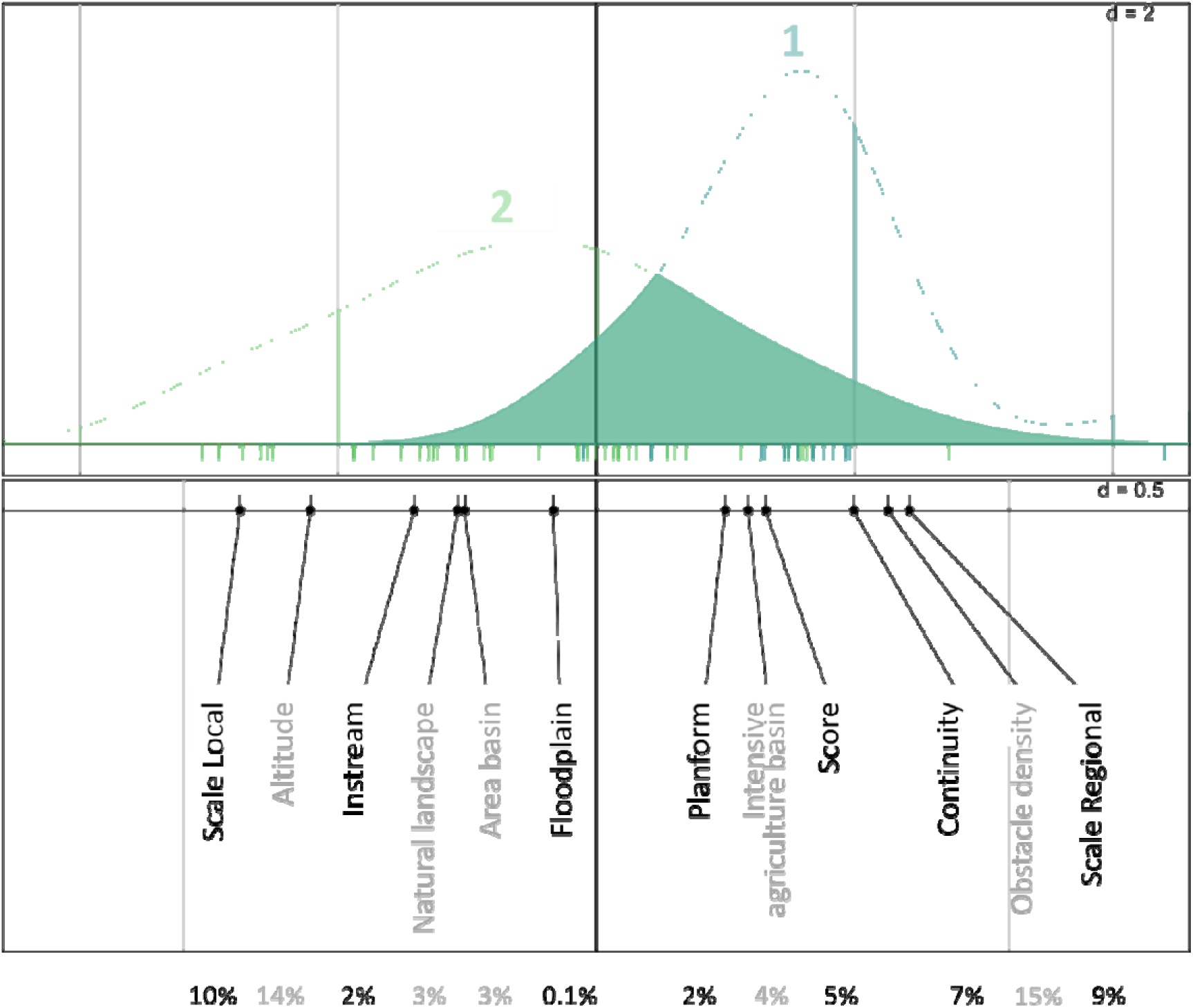
Characterisation of the clusters by environmental variables and restoration measure types for macroinvertebrates. Above: representation of the Between-Group Analysis (BGA), with cluster 2 on the left, in green, and cluster 1 on the right, in blue. The vertical lines under the density curves represents the position of the sites on the axis. Below: representation of the correlation with the environmental variables (grey) and restoration measure types (black). The more the variable (or the modality) is on the right of the axis, the more it is correlated with cluster 1 and the more the variable (or the modality) is on the left, the more it is correlated with cluster 2. The percentages represent the absolute contribution of each variable to the explained variation of the BGA.

Cluster 1, which is characterised by a predominance of affirmative responses, is typified by sites that have undergone restoration projects at a regional, targeted scale. Furthermore, cluster 1 incorporates sites where the restoration project comprised multiple measures (a high "Score"), including measures that enhanced river continuity – and to a lesser extent planform morphology. Cluster 1 is distinguished by its location in basins characterised by intensive agricultural practices. These sites exhibit a high density of obstacles, which consequently results in a high level of fragmentation in the reaches. Conversely, sites classified within cluster 2, i.e. sites exhibiting no response or negative response to restoration, are distinguished by their elevated altitude, integration within a more pristine landscape, expansive river basins, and a project focused on a local scale, with measures enhancing instream morphology.

## Discussion

### 1. Whether river restoration outcome is positive or not depends on environmental context and restoration characteristics

#### a. Only one quarter of the restoration projects led to positive outcomes

The growing body of literature that aims at assessing the ecological outcomes of river restoration shows contrasting and variable results, and pinpoints a need for an integrative method to obtain a comprehensive assessment of restoration outcomes linked to both the environmental context and restoration characteristics. In this multi-metric study on macroinvertebrates, we showed that taxonomic and functional diversity and richness, as well as sensitivity and dispersal metrics, tend to have the same aggregated response over 3 to 6 years after restoration. Two groups of sites differing in the direction of their responses to restoration were observed: positive or neutral/negative outcome. This clear dichotomy is quite unexpected, as the studies assessing river restoration outcomes show, using different metrics, very variable restoration outcomes, going from negative to positive effects(see Introduction, (Palmer et al. 2010; Schmutz et al. 2014). In light of the heterogeneity of the sites encompassed in our analysis, we sought to elucidate this dichotomy by environmental context (i.e. agricultural land use, fragmentation (i.e. density of obstacles)) and restoration measure (i.e. river continuity and planform morphology, targeted scale of the project) characteristics.

#### b. Environmental context and restoration characteristics equally influence restoration outcomes

The findings of this study demonstrate that the observed variation in restoration outcomes can be partially attributed to both environmental context and restoration characteristics. Each of these factors contributes equally to the variability that is explained. For instance, higher altitude sites have been observed to exhibit a tendency towards neutral or negative ecological responses to restoration initiatives. This phenomenon can be attributed to the reduced taxonomic richness of macroinvertebrates with altitude (Jacobsen 2003; Suren 1994), resulting in smaller species pools and, consequently, diminished opportunities for richness to respond positively to restoration measures. A further hypothesis to explain the diminished response to the restoration of sites at higher altitudes is that rivers at higher elevations may be in a more natural state prior to restoration than those at lower elevations. This would result in a lower response ratio.

The results obtained demonstrate that macroinvertebrate communities exhibit a robust response to degradation, particularly in the context of fragmented and agriculturally impacted environments. The outcomes of the restoration projects were influenced by pressures related to land use at the basin level, with stronger responses observed in sites exhibiting high levels of intensive agriculture (also shown for fish, Kail et al., 2015; Lorenz et al., 2013). It is evident that the observed species richness is a direct consequence of the pre-restoration species richness, which is notably lower in areas exhibiting high agricultural intensity when compared to those demonstrating low agricultural intensity. This phenomenon has also been observed in the context of forestry, as evidenced by Egler et al. (2012). This has the effect of accentuating the difference in diversity between communities that were initially poor and those that were initially rich, before and after restoration.

The findings of this study demonstrate that ambitious restoration initiatives can yield favourable ecological outcomes. Indeed, the restoration of sites that have been successful is characterised by the implementation of a regional-scale target. This finding aligns with the extant literature on macroinvertebrates, which documents that restoration outcomes are often suboptimal in stretches that are restored over less than 2 km (Haase et al., 2013; Hering et al., 2015). However, Hering et al. (2015) highlighted that the length of the restored section is not sufficient in itself to explain restoration effects in biota. In this particular instance, the implementation of restoration initiatives encompassing comprehensive measures has been demonstrated to elicit more pronounced positive responses. Specifically, measures directed towards enhancing river continuity and planform morphology have been observed to yield more favourable ecological outcomes in comparison to those aimed at improving instream morphology. This finding is in contrast with the conclusions of other studies, which indicated that instream measures resulted in more successful restorations (Al-Zankana et al. 2020; Kail et al. 2015). The present study demonstrates that the implementation of restoration projects comprising a variety of measure types results in more favourable ecological outcomes (i.e. a higher ‘restoration score’). This assertion challenges the conventional approach of selecting one measure type over another.

Furthermore, it was highlighted that the group of sites exhibiting positive outcomes is associated with both measures pertaining to the continuity of the river and a higher level of fragmentation of the reach (i.e. obstacle density). This phenomenon may be attributable to two distinct yet not yet disentangled effects, which are not mutually exclusive, as one may be applicable to some sites while the other may be applicable to other sites. Firstly, the restoration of river continuity has been demonstrated to enhance the colonization of macroinvertebrates by avoiding less optimal conditions in backwater dams and by facilitating dispersal. As, Li et al. (2016) and England et al. (2021) have demonstrated, dispersal is a pivotal factor in the success of macroinvertebrate restoration. Secondly, the outcomes observed in terms of ecological systems may be attributable to the proximity and composition of source populations in the vicinity of restored sites (Tonkin et al. 2014). In this case, the hypothesis is that higher levels of fragmentation in the restored sites landscape may have created more diverse habitats (Frainer et al. 2018; Verdonschot et al. 2016) in turn enhancing more diverse macroinvertebrate communities and enabling higher recolonization of the restored sites.

In conclusion, it is demonstrated that the environmental aspect is to be given equal consideration to the type of restoration implemented when restoration projects are conceived. This is due to the fact that both play an equal part in explaining the variability observed in ecological outcomes.

### 2. Insights from multi-metric analysis for a comprehensive assessment of river restoration outcomes

The present study further demonstrates how the multi-metric analysis of river restoration outcomes, and in particular the use of complementary taxonomic and functional metrics, allows a comprehensive assessment of river restoration. Firstly, it was demonstrated that restorations which were associated with strong increases in taxonomic richness were accompanied by moderate increases in trophic-level functional diversity, low increases in overall functional diversity, and even negative functional evenness. This discrepancy between taxonomic and functional diversity metrics may be indicative of the necessity for a longer period for functional diversity metrics to reach their optimal response to restoration (Baker et al. 2021; Nguyen et al. 2024; Rideout et al. 2022). This delay between taxonomic and functional responses can be attributed to two competing hypotheses: functional redundancy or keystone species effects (Naeem 1998; Ali 2023; Boulton et al. 2008; Schulze et al. 2019). In accordance with the functional redundancy hypothesis, the colonisation of a restored reach by new species results in an increase in taxonomic richness without concomitant increase in functional richness, on the grounds that the functions of the new species are already present in the community. In accordance with the keystone species hypothesis, the restored reach must accumulate a substantial number of species before it can acquire new functions, as these functions are ensured by only a few rare, key species. It can be reasonably assumed that the first hypothesis of functional redundancy can most likely explain the observed gaps between taxonomic and functional diversities, if it is considered that the restored reaches already have relatively rich communities from 3 to 6 years after restoration. Secondly, it was demonstrated that the utilisation of trait-based metrics pertaining to sensitivity and dispersal ability facilitated the refinement of comprehension regarding restoration outcomes. For instance, it was determined that stenothermic species and those with a high capacity for dispersal may derive the greatest benefit from restoration processes; conversely, the majority of sensitive taxa (EPT) exhibited no recovery. This finding indicates that, within the designated study sites, the restoration process has yielded partial success in terms of ecosystem function, as evidenced by the recovery of species that exhibit high levels of natural tolerance and resilience, from three to six years following the initial restoration. It is evident that no study has hitherto been conducted that has demonstrated these simultaneous responses of dispersal capacity, stenothermy and EPT species richness within a single integrative framework.

Despite the fact that these three metrics have been studied separately on a number of occasions (for example, 80% of studies meta-analysed by (Al-Zankana et al. 2020) do not demonstrate an increase in EPT richness). These novel insights into the varied ecological responses to river restoration were obtained through the integration of taxonomic and functional metrics, as revealed by our multi-metric assessment. In order to further explore the mechanisms that drive restoration outcomes, one could, for instance, test specific hypotheses concerning each ecological response by analysing their dynamics over time following restoration.

### 3. Future perspectives on river restoration assessment

The results obtained from the analysis of multi-metric and multi-contextual datasets through the implementation of clustering and descriptive analyses have indicated the emergence of patterns in river restoration outcomes that are of interest. However, the present analyses are restricted to a limited post-restoration period (3 to 6 years), thus capturing the ecological status of the restored sites on a limited time frame. In contrast, extant literature demonstrates that there can be temporal variability in biodiversity responses to restoration. Indeed, the temporal evolution of the ecological responses may vary according to the nature of the considered metrics or the type of restoration measure applied. Indeed, the temporal aspect constitutes an alternative explanation for the variability in restoration outcomes that was not addressed in this study. A number of studies have been conducted which present repeated monitoring data (Schmutz et al. 2014). These studies demonstrate that ecological responses are highly dynamic over time. The dynamic nature of ecological responses is evident in a number of ways. Firstly, they are dependent on the taxa under study (Lepori et al. 2005).

Secondly, they are dependent on the proximity of the source population of potential colonisers (Stoll et al. 2016; Tonkin et al. 2014). Thirdly, they are dependent on the duration of the life cycle. Consequently, it would be a worthwhile endeavour to explore the temporal pattern of the clustering over time. A period of 3 to 6 years after restoration allows for the observation of differences in responses of sites; however, it is expected that in some sites, ecological metrics and restoration measures require additional time to demonstrate stronger ecological responses to restoration.

Furthermore, it is evident that the two clusters elucidated a mere 7.6% of the aggregate variance in ecological responses. It is evident that, despite the capacity to derive reliable conclusions from the results (as evidenced by the permutation test of the BGA achieving significance and the spuriousness index attaining a remarkably low value), the influence of other factors on the association with the clusters remains a salient factor. In order to enhance comprehension of the ecological responses to river restoration, it is imperative to explore and assess the influence of additional factors. These factors may include the geographical location of the river continuum (Manfrin et al. 2019), water quantity and quality (Brettschneider et al. 2023; Jähnig et al. 2010; Palmer et al. 2010), legacy effects (Chen and Olden 2020)), or even factors associated with the nation in which the restoration project was implemented. Indeed, as demonstrated in Appendix 4 (Figure S1), significant disparities in the distribution of measure types were observed between countries. The selection of appropriate measurement types and their alignment with the environmental context of the restoration site may be contingent on the restoration policies that differ between countries.

### 4. Conclusion

The clustering and descriptive analyses employed in this study to investigate ecological metric responses to restoration enabled the following: i) the characterisation of predominant signals by identifying sites where macroinvertebrate ecological metrics exhibited analogous responses and ii) the determination of the environmental context and restoration characteristics of the sites belonging to the various clusters. The present study proposes an integrative approach that responds to the need for novel methods to assess river restoration outcomes using multi-dimensional biodiversity metrics and multi-contextual data. In view of the findings, it is recommended that this analysis be adopted more broadly (Al-Zankana et al. 2020; Palmer et al. 2010; Sinclair et al. 2023; Stoll et al. 2016). From the standpoint of a water manager, the present study demonstrates the significance of site selection in the planning of restoration measures, as restoration effectiveness or success rate can vary considerably among sites, rivers and basins. This study is therefore part of the recent strategic restoration and conservation planning movement for river restoration (England et al. 2021; Weber et al. 2018). It is demonstrated that the implementation of restoration initiatives can yield more favourable ecological outcomes (encompassing both taxonomic and functional metrics) for macroinvertebrate communities, provided that the restoration is ambitious, encompassing a substantial targeted scale and incorporating a variety of metrics, and is executed in locations that are degraded and fragmented.

## Acknowledgments

This work is part of the COSAR project, joint ERA-Net Biodiversa+ / WaterJPI 2021 BiodivRestore Call, and has been funded through French ANR (ANR-21-BIRE-0001 and ANR-21-BIRE-0002), German DFG (491738349), Dutch MINLNV (BO-43-222-012) and Swiss EAWAG. Blandine Charrat has also been supported by the Graduate School H2O’Lyon (ANR-17-EURE-0018) of Université de Lyon (UdL), within the program "France 2030” operated by the French National Research Agency (ANR).

## Author Contributions

Blandine Charrat, Jérémy Piffady and Céline Le Pichon conceptualized the study. Mathieu Floury curated the data. Blandine Charrat analysed and visualized the data and wrote the manuscript. All authors contributed to reviewing and editing the manuscript.

