## Appendices for "Towards a multi-metric assessment of river restoration ecological outcomes across projects and contexts"

**Manuscript type:** Research manuscript

[1]: INRAE, UR RiverLy, EcoFlowS, Villeurbanne, France

[2]: University of Applied Sciences Trier, Environmental Campus Birkenfeld, Hoppstädten-Weiersbach, Germany

[3]: University of Paris-Saclay, INRAE, UR HYCAR, Antony, France

[4]: Faculty of Biology, University of Duisburg-Essen, Essen, Germany

[5]: Wageningen Environmental Research, Wageningen University and Research, Droevendaalsesteeg 3, 6708 PB, Wageningen, The Netherlands

[6]: Eawag, Swiss Federal Institute of Aquatic Science and Technology, Surface Waters – Research and Management, Kastanienbaum, Switzerland

**Appendix S1: Description of the traits considered for the functional metrics**

Table S1: List of traits and modalities from Tachet et al. (2010)[(1)] used to compute the functional indices. Description of the traits and modalities for habitat, dispersal and mobility, organism size and form, life history and reproduction and feeding groups for the trait groups included in functional metrics calculation.

| Trait group | Trait | Modalities |
|---|---|---|
| Habitat | Substrate preference | boulders to pebbles |
| | | gravel |
| | | sand |
| | | silt |
| | | macrophytes |
| | | microphytes |
| | | twigs/roots |
| | | organic detritus/litter |
| | | mud |
| | Current velocity | Null |
| | | slow |
| | | medium |
| | | fast |
| | Temperature | psychrophylic |
| | | thermophilic |
| | | eurythermic |
| Dispersal and mobility | Dispersal mode | aquatic passive |
| | | aquatic active |
| | | aerial passive |
| | | aerial active |
| | Locomotion | flier |
| | | surface swimmer |
| | | full water swimmer |
| | | crawler |
| | | burrower |
| | | interstitial |
| | | temporarily attached |
| | | permanently attached |
| Organism size and form | Maximal body size | ≤ 0.25 cm |
| | | > 0.25-0.5 cm |
| | | > 0.5-1 |
| | | > 1-2 cm |
| | | > 2-4 cm |
| | | > 4-8 cm |
| | | > 8 cm |
| Life history and reprodution | Life cycle duration | ≤ 1 year |
| | | > 1 year |

| | | |
|---|---|---|
| | Number of cycles per year | < 1 |
| | | 1 |
| | | >1 |
| | Aquatic stages | egg |
| | | larva |
| | | nymph |
| | | adult |
| | Reproduction | ovovivipary |
| | | isolated eggs, free |
| | | isolated eggs, cemented |
| | | clutches, cemented or fixed |
| | | clutches, free |
| | | clutches, in vegetation |
| | | clutches, terrestrial |
| | | asexual reproduction |
| Feeding group | Feeding habit | absorber |
| | | deposit feeder |
| | | shredder |
| | | scraper |
| | | filter-feeder |
| | | piercer |
| | | predator |
| | | parasite |

(1) Tachet H, Richoux P, Bournaud M, Usseglio-Polatera P (2010) Invertébrés D'eau Douce: Systématique, Biologie, Écologie. CNRS Editions, Paris.

**Appendix S2: Determination of the optimal number of clusters according to the average silhouettes method**

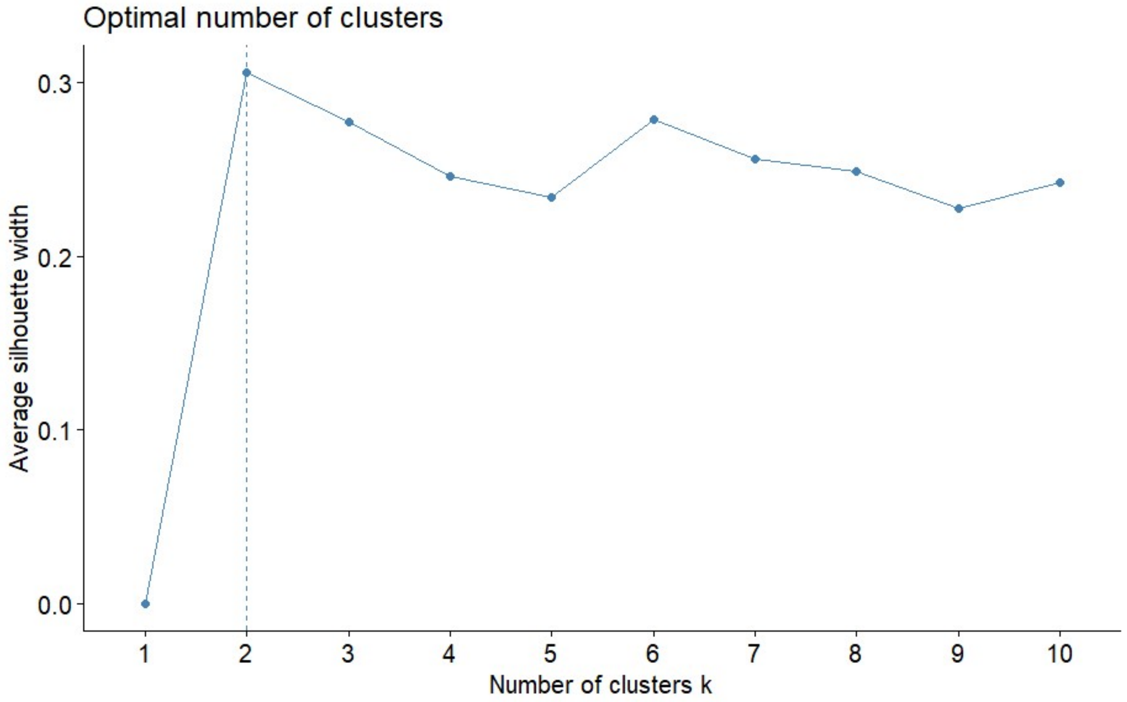


Figure S1: Representation of the average silhouettes width according to the number of clusters. The dotted line indicates that 2 clusters lead to the highest average silhouettes width, and thus is the optimal number of clusters.

**Appendix S3: Hill-Smith analysis representation**

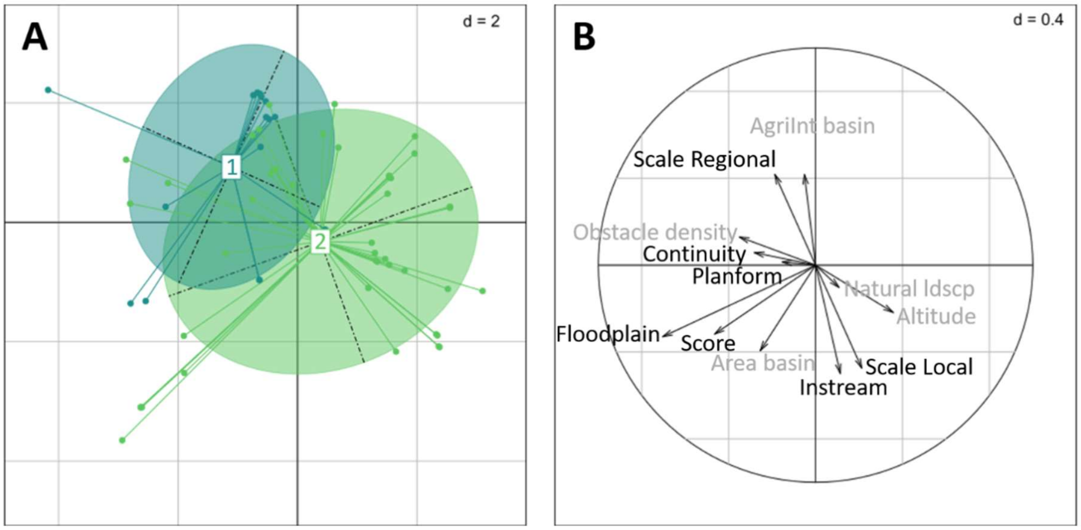


Figure S1: Representation and correlation circle of the Hill-Smith analysis led on the restoration characteristics and environmental context variables. A: representation of the 62 restored sites in the space of the Hill-Smith analysis. The identity of the cluster is used as an illustrative variable, showing in blue sites that belong to cluster 1 and in green sites that belong to cluster 2. B: correlation circle of the Hill-Smith analysis. In black are the variables linked to the characteristics of the restoration project and in grey are the variables linked to the environment.

**Appendix S4: Different measure types across the three studied countries**

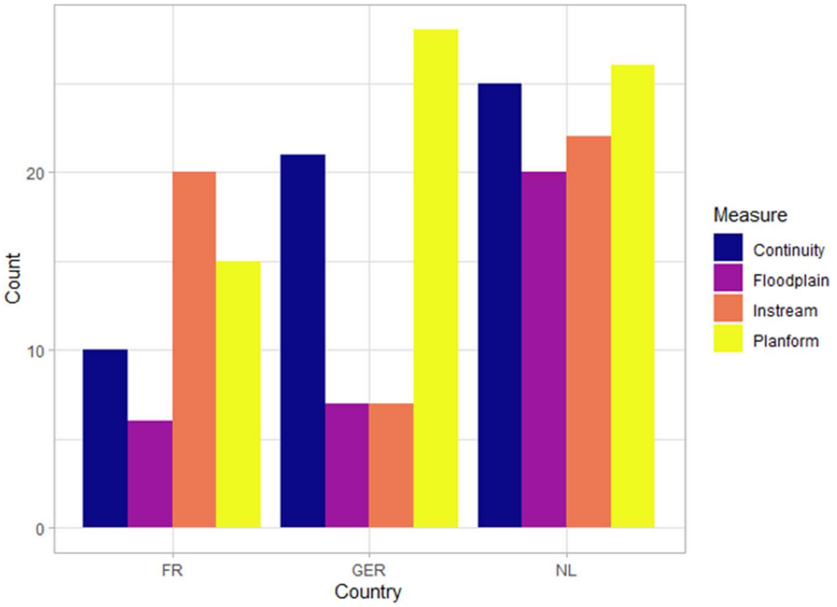


Figure S1: Distribution of measures between countries. FR = France, GER = Germany, NL = The Netherlands. Continuity = restoration measures that aim at improving river continuity, Floodplain = restoration measures that aim at improving floodplain morphology, Instream = restoration measures that aim at improving instream morphology, Planform = restoration measures that aim at improving planform morphology.
